# Signal peptides as potential structural modulators of human splice isoforms

**DOI:** 10.1101/2025.05.27.656295

**Authors:** Tomer Tsaban, Ora Schueler-Furman

## Abstract

Signal peptides (SPs) are short protein segments responsible for protein localization, typically cleaved from mature proteins and rapidly degraded after fulfilling their targeting function. Beyond localization however, these sequentially and structurally diverse elements may play additional roles. We explore how alternative splicing potentially creates structural contexts where SPs become integral components of folded domains. Employing AlphaFold and additional computational approaches, we examined a known functional immune protein splice isoform of human SLAMF6 that retains biological activity, despite forming a truncated domain that lacks the central elements of the canonical interaction interface. We revealed striking characteristic similarities between the SLAMF6 SP and the absent segment, indicating its ability to complement and stabilize the isoform domain. An *in silico* screen of 235,000 reported expressed human isoforms identified several dozen additional candidates with potential SP complementation, previously dismissed as modeling artifacts. Notably, immunoglobulin and carbonic anhydrase domains show particular enrichment among these candidates. SPs are commonly regarded as dispensable elements that can be altered or eliminated when investigating proteins and their structures. Our research proposes an alternative perspective wherein SPs might perform integral roles in stabilizing their source proteins, or others. These findings extend the growing body of evidence for moonlighting SPs, suggesting that we have only begun to uncover their true functional scope. Specifically, SPs emerge as unique modulatory elements essential for understanding the structural and functional behavior of protein splice isoforms.

## Introduction

Signal peptides (SPs) are N-terminal protein segments who play a major role in protein localization. Most transmembrane and secreted proteins require a signal peptide in order to be directed into the ER lumen, where they are folded, and from which they are then trafficked outside the cell. Usually, the signal peptide is recognized by a signal recognition particle complex, and is cleaved off to leave a mature protein at the target site (e.g., the cell membrane). For most SPs, degradation follows quickly after cleavage (1),(2),(3). SPs are highly variable in sequence (4),(5),(6). They are usually composed of 3 regions: First, a positively charged terminus; second, a hydrophobic α-helix; and third, a hydrophobic/uncharged β-strand that often defines the cleavage recognition site (2). While very common, these secondary structures are affected by the environment which may induce a secondary structure shift (1),(7). This versatile nature suggests additional potential functions for SP segments, *i.e.,* SP moonlighting. Indeed, since their discovery, evidence has accumulated to suggest that SPs may have structural roles in addition to localization (3),(2), including (but not limited to) binding lipids (8),(9),(10), oligomeric state modulation (11), and even forming peptide-protein interactions after their cleavage (12),(13),(14). Some evidence exists for SPs who affect the folding process of the same protein that they localize (15),(16) and in viruses, SPs have been shown to have integral structural and functional roles, including stabilization of mature proteins (17),(18),(19).

However, SPs who play integral structural roles in mature human proteins have not yet been reported. Such an additional role for SPs could be revealed in alternative splicing isoforms - where elements are often removed, truncated or shifted in order, creating new structural contexts for SPs to take part in. Alternative splicing is used in higher eukaryotes to increase the functional variability of genes, in response to changing cellular settings. Such splicing leads to inclusion/skipping of specific exons, and sometimes to changes of protein sequence due to frameshifts. Alternative splicing may also affect the SP sequence. Recently, Hajaj *et al.* demonstrated the expression and function of a naturally occurring splice isoform of the gene SLAMF6, “variant-3” (20), which we will call here isoform3. While the canonical, most frequent variant of the protein is a well described T-cell inhibitor (21),(22), isoform3 was found to be a T-cell agonist (20). This remarkable system of a complete switch (on/off) based on two products of the same gene, is possible through alternative splicing. Surprisingly, the two isoforms are identical in sequence, except for a 58 amino acid segment (G17-I65) that is missing from isoform3, effectively removing a significant part of the first SLAMF6 domain, including a central part of the interface by which SLAMF6 homodimerizes for signal transduction. Interestingly, the deletion also removes the cleavage site of the SP at residues 20-21 (23),(24), leading us to believe that cleavage might be reduced or even abolished. Still, both SLAMF6 canonical and isoform3 are functional and expressed on the membrane (20),(22), indicating a functional SP. Aside from its role in localization, we hypothesized that it could structurally compensate for the missing segment, thereby stabilize isoform3 and also contribute to an altered dimerization interface.

Protein isoforms remain challenging to study both experimentally (25),(26),(27) as well as computationally (28),(29),(30),(31),(32) even today. For many isoforms, their translation into protein and their biological relevance has not yet been demonstrated. The standing challenge of differentiating meaningful splice isoforms from sequencing noise (33),(34),(35),(36), lead to the design of comprehensive resources such as the CHESS curated isoform database (37) and the MANE consensus (canonical) isoform set (38). Structural modeling of these isoforms can be challenging as well.

Computationally, state-of-the-art structural modeling methods such as AlphaFold2 (AF) have been thoroughly validated (39, 40). However, many rely heavily on multiple sequence alignment (MSA) to draw co-evolutionary restraints that define the protein structure (41). Splice isoforms are less represented in sequence databases (36) and may be confounding for MSA generating algorithms (42),(43), given their existence mainly in higher life forms and often high partial similarity to their canonical counterparts. This makes it challenging to identify isoform-specific features due to obvious bias towards the canonical structure. Such bias was recently observed where AF “ignores” subtle co-evolutionary signals in fold switching proteins and tends to only predict the canonical fold (44),(45, 46). Zhang *et al.* have shown that AF (and other similar approaches) tend to model isoforms as an “incomplete”, partial subset of the canonical fold, rather than predicting a conformational, or even a fold change, even when the incomplete structure seems biophysically less likely (47).

But what if some isoforms actually form incomplete folds? These structures could be stabilized by alternative elements, in a similar fashion to peptide-protein interactions “mimicking” or “recreating” folded cores (48),(49),(50). In particular, could SPs complement such predicted partial folds? A promising approach to distinguish “real” partial folds from modeling artifacts (due to MSA bias or fold memorization) is based on large protein language models (pLMs) such as ESM (51) which were not trained on structural data or MSA, but rather on massive datasets one sequence at a time, in an unsupervised manner. Relationships between local elements can be extracted from these ESM logits (embeddings) in the form of a categorical Jacobian (47).

In this study, we sought to characterize structural and functional contributions of SP in splicing isoforms, using a variety of complementary computational methods. We first studied the structure of SLAMF6 isoform3 and characterized the potential role of its SP in stabilizing a domain and complementing its interface. Our analysis suggests specific structural alterations that could help explain domain conservation on the one hand, and functional diversity on the other hand (compared to the canonical protein). Following the SLAMF6 example, we used a similar computational strategy to detect several dozen protein isoforms where SP complementation may occur. We provide sequence-based, co-evolutionary, and mass spectrometry (MS) evidence to support this hypothesis. This work suggests a new potential mechanism by which SPs stabilize protein isoform domains and interfaces, and may shed light into nature’s ability to diversify the protein repertoire by repurposing existing elements.

## Results

### Structural basis for SLAMF6 signaling via homophilic interaction

SLAMF6 is a member of the signaling lymphocytic activation molecule (SLAM) family. SLAM receptors are widely expressed on immune cell surfaces and interact via homodimerization, either homotypic (2 molecules on the same cell) or heterotypic (2 molecules, one from each cell). Structurally, SLAMF6 contains two extracellular Immunoglobulin like domains: an N-terminal variable (V) domain (**Figure 1A,B**), followed by a linker and a constant (C) domain. Separated by a transmembrane domain, the intracellular regions regulate signal transduction to the cell via phosphorylation. The extracellular interaction between two SLAMF6 molecules is formed by an interface between the front faces of the N-terminal V domains. These two-layered β-sheet Ig-like domains consist of 6 front β-strands and 3 at the back. The solved crystal structure of the canonical isoform of SLAMF6 reveals that the homophilic interface is predominantly formed by C, C’ and F strands (24) (**Figure 1B**). Alanine mutations in these strands (at positions F52, L56, E59, F64) cause interface destabilization and even misfolding (F64), while other residues (I53, W55, I65) stabilize the domain core and are highly conserved (24) (**Figure 1A**). This V domain-mediated homophilic interaction is at the basis of human T-cell inhibition by SLAMF6. In contrast to this canonical isoform, a naturally occurring SLAMF6 splice isoform, isoform3, has been reported to have an agonistic effect and activate T-cells (20). The isoform shows a unique splice pattern, in which half of the V Ig-like domain is removed (**Figure 1A**). Of special importance, strands C and C’ are missing (**Figure 1B**).

**Figure 1:**
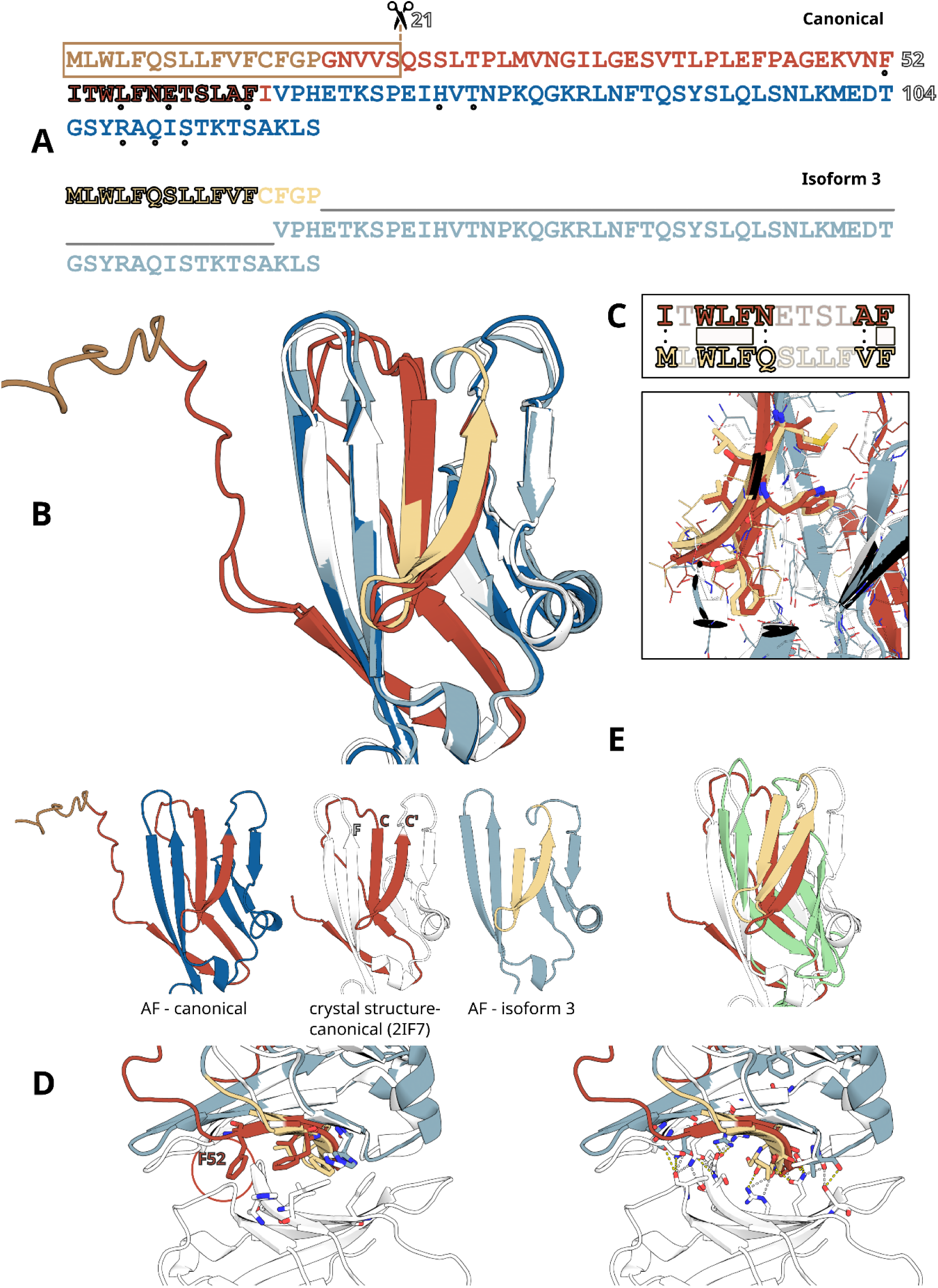
The signal peptide of SLAMF6 isoform3 complements the missing interface segment. **(A)** Sequence of SLAMF6 V-domain of the canonical isoform (CHS.3843.5; top, dark colors) and isoform3 (CHS.3843.1; bottom, light colors). The canonical signal peptide (SP) is highlighted by a box. Complementing and complemented residues are outlined in black, and shown in detail in panel **C**. Coloring scheme: SP: brown/wheat; common sequence: blue; residues in the canonical protein that are missing in isoform3: dark red. Hotspot residues are marked with black dots. **(B)** Superimposed models of N-terminal V-domain of SLAMF6 demonstrate overlap of isoform SP with missing segment. Coloring scheme for AF models as in **A.** Crystal structure of the canonical isoform (PDB ID 2if7 (24)): white. The interface involved in dimerization is shown in the front. **(C)** Sequence complementarity of the isoform3 SP with missing (complemented) interface segment, recovery of intra-domain stabilizing interactions and overlap of the WLF-F motif. **(D)** Isoform3 model superimposed with the crystal structure homodimer interface suggests recapitulation of crucial interactions: right panel - recovery of hydrogen bond network; left panel - recovery of hydrophobic interactions, except for F52 (marked by a circle). **(E)** Isoform3 model generated by Rosetta *ab-initio*, a traditional, pre-deep learning method (pale green), demonstrates similar complementation by the SP (see **Supplementary Figure S2** for corresponding I-TASSER models).

### The signal peptide of isoform3 likely complements its truncated domain

In the canonical protein, the SP is cleaved around positions 21, resulting in a mature protein that begins with the V domain at residue 22 (**Figure 1A**) (23),(24). Isoform3 lacks residues 17-65, including the cleavage site, but is otherwise identical to the canonical protein (**Figure 1A**). Given the removal of the last 5 residues of the SP, we assumed that its cleavage may be altered and perhaps abolished, without disrupting the localization signal (as supported by the extracellular detection of this isoform in Hajaj *et al.* (20)). Under this assumption, we inspected the sequence of isoform3 in alignment to the canonical protein. Surprisingly, the retained part of the SP in isoform3 contains a motif nearly identical to the one removed at the center of the interaction: residues 1-ML**WLF**QSLLFV**F**-12 of the SP are well aligned to strands C and C’ of the canonical protein (53-IT**WLF**NETSLA**F**-64) and specifically key residues 55-WLF-57 and F64 are preserved (**Figure 1C**).

The sequentially aligned segment has a propensity to form an α-helix followed by a β-strand in the canonical context (2),(1). However, in the isoform context, it is likely to “shift”, and form beta-strands only, emphasizing its structural compatibility with the missing β-hairpin. This is supported by evidence in the literature of SP helix-to-strand transitions upon changes in the environment (1),(7). The sequence similarity, secondary structure compatibility and expected alteration of the SP cleavage site, led us to believe that the SP of SLAMF6 isoform3 is likely to form a β-hairpin structure, and complement the missing C-C’ strands at the interaction interface. Reassuringly, AF models of isoform3 support this hypothesis, “folding” the SP in a way that complements the V domain (**Figure 1B**), thereby precisely recovering the 55-WLF-57-F64 side chain conformations (**Figure 1C**). Superposition of the isoform model onto the crystal structure dimer reveals recovery of most of the crucial interactions of the interface (polar as well as hydrophobic, **Figure 1D**). However, the dimerization interface differs from the canonical interface, in particular, the interaction formed by F52 which is crucial for dimerization, is not recovered in the isoform (**Figure 1D**, circle). We believe this similar but partial interface could account for the change in dimerization described previously (20).

### Co-evolutionary analysis supports the structural complementation of the SP in isoform3

While the AF predictions support our hypothesis that the SP could stabilize the truncated V domain in SLAMF6 isoform3, they could be biased. Protein isoforms are known to be challenging targets for computational structure prediction; Their sequence is often similar to the canonical sequence and they are less conserved throughout evolution, biasing MSA-based predictions towards the canonical proteins. Additionally, they are often harder to study experimentally, leading to underrepresentation in structural databases like the Protein Data Bank (PDB) (32). These biases are especially strong with homology-based, template-based, or coevolution-based methods such as AF (which was trained on homologs, and derives contacts based on co-evolution signals detected in MSAs). Indeed, a recent study that examined structural models of 18 protein isoforms deemed these models as unlikely based on exposed hydrophobic residues, concluding they are incomplete copies of canonical domains as a result of the aforementioned biases (47). Thus, in principle, the isoform3 model generated by AF could result from such a bias, while in fact isoform3 adopts a different conformation, or even unfolds (**Figure 2A**). Reassuringly however, we provide two (non-exhaustive) “counter examples”: isoform models of MYG and CAS9 examined by that same study (47) may be well explained in the context of larger protein complexes and thus fold as predicted (**Supplementary Figure S1**). We therefore believe AF models of isoforms are not a-priori wrong for being partially “copied”. Furthermore, we were able to generate highly similar models for isoform3 using other, pre-deep learning methods that are less prone to the described biases: Rosetta *ab-initio* (that does not use neither structural templates nor MSA-based contact information) (52) (**Figure 1E**) and I-TASSER (53) (**Supplementary Figure S2**). This supports the credibility of the SP complementation model of isoform3.

**Figure 2:**
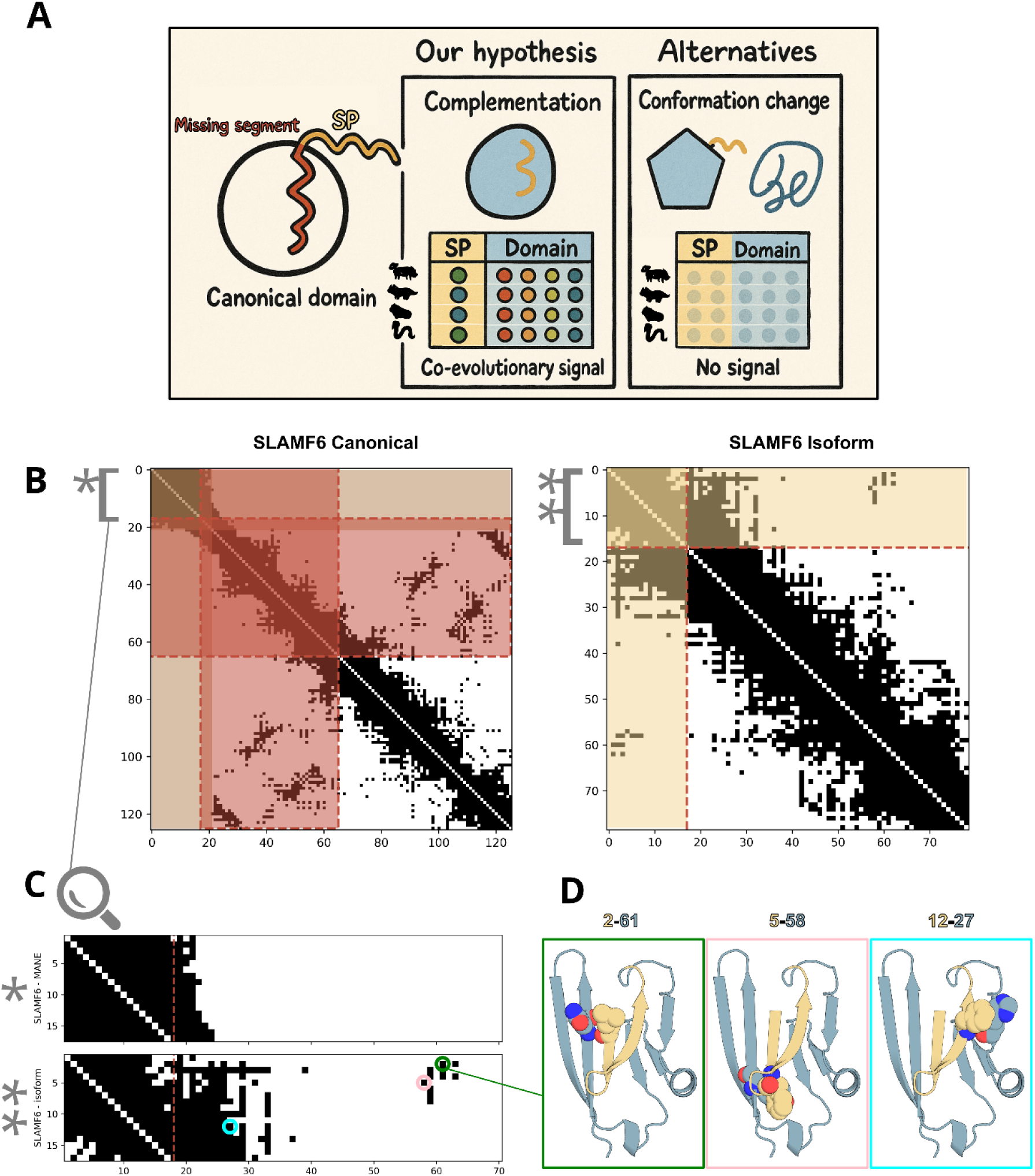
Co-evolutionary analysis supports SP complementation in SLAMF6 isoform3. **(A)** Possible folding alternatives for SLAMF6 isoform with regards to the missing segment (dark red) and SP (wheat). **(B-E)** Co-evolutionary analysis of the SLAMF6 canonical (MANE consensus isoform) and isoform3 sequences using ESM-2-based categorical Jacobians. **(B)** Predicted categorical Jacobian contact maps of canonical isoform (right) and isoform3 (left). The SP area is marked brown and wheat respectively; the missing segment is marked dark red. **(C)** Zoom-in into the contact map segment of the SP-domain interaction reveals an “emerging” signal in isoform3 that is missing in the canonical isoform. **(D)** Structural visualization of selected residue pairs in isoform3 on the AF model structure. Outline colors correspond to circles in the contact map.

We speculated that an independent analysis of co-evolving residues could further distinguish probable SP complementation from a structural modeling artifact. We hypothesize that if indeed SP complementation occurs, a co-evolutionary signal “connecting” the SP region and following domain in isoform3 should be detected. Otherwise, if isoform3 changes conformation without the involvement of the SP or completely unfolds, no such co-evolutionary signal should be found. The latter would suggest that the AF isoform3 model is predicted because AF is unable to predict conformational changes, and has simply copied the canonical structure (**Figure 2A**). To test this, we inspected co-evolution signals, as detected in the categorical Jacobian (47) of ESM-2, an MSA-independent language model (51). For the canonical SLAMF6, the Jacobian reveals a strong co-evolutionary signal within the SP region (SP to itself), but no signal between the SP and the following domains. For isoform3, however, new residue pairs are predicted to be associated (and potentially co-evolved), suggesting new interactions between the SP region and the domain that follows (**Figure 2B**). Specifically, the Jacobian predicts interactions between residue pairs 2-61, 5-58 and 12-27, strongly constraining the structure of isoform3 to the suggested AF model, with the SP complementing the missing segment (**Figure 2C,D**). These results strongly support SP complementation of the isoform3 V domain, in accordance with the corresponding AF models. We thus conclude that the AF models of isoform3 are unlikely to be solely based on biased fold memorization.

In summary, the SP complementation model of SLAMF6 isoform3 is supported by several levels of evidence - sequence compatibility and motif similarity, independent structural models and co-evolutionary analysis. While not fully independent, these lines of evidence are not bound to be in agreement. Hence, we find this cluster of complementing results in favor of a true signal, rather than method-related artifact. Encouraged by this, we hypothesized that signal peptides may have a broader structural role within mature protein domains, and specifically for protein isoforms.

### Large scale computational screen of human splice isoform SPs

How recurrent is this SP-mediated complementation in splicing isoforms? To evaluate the frequency of the suggested phenomenon observed for SLAMF6 isoform3, we designed a computational pipeline (**Figure 3**). We restricted our analysis to likely biologically relevant human protein isoforms, as defined in the curated CHESS database (37),(54), and defined the “canonical” isoforms based on the MANE annotation of consensus isoforms (38) (237,275 CHESS isoforms, of which 33,851 are annotated as MANE canonicals, and the remaining 203,424 are alternative isoforms).

**Figure 3:**
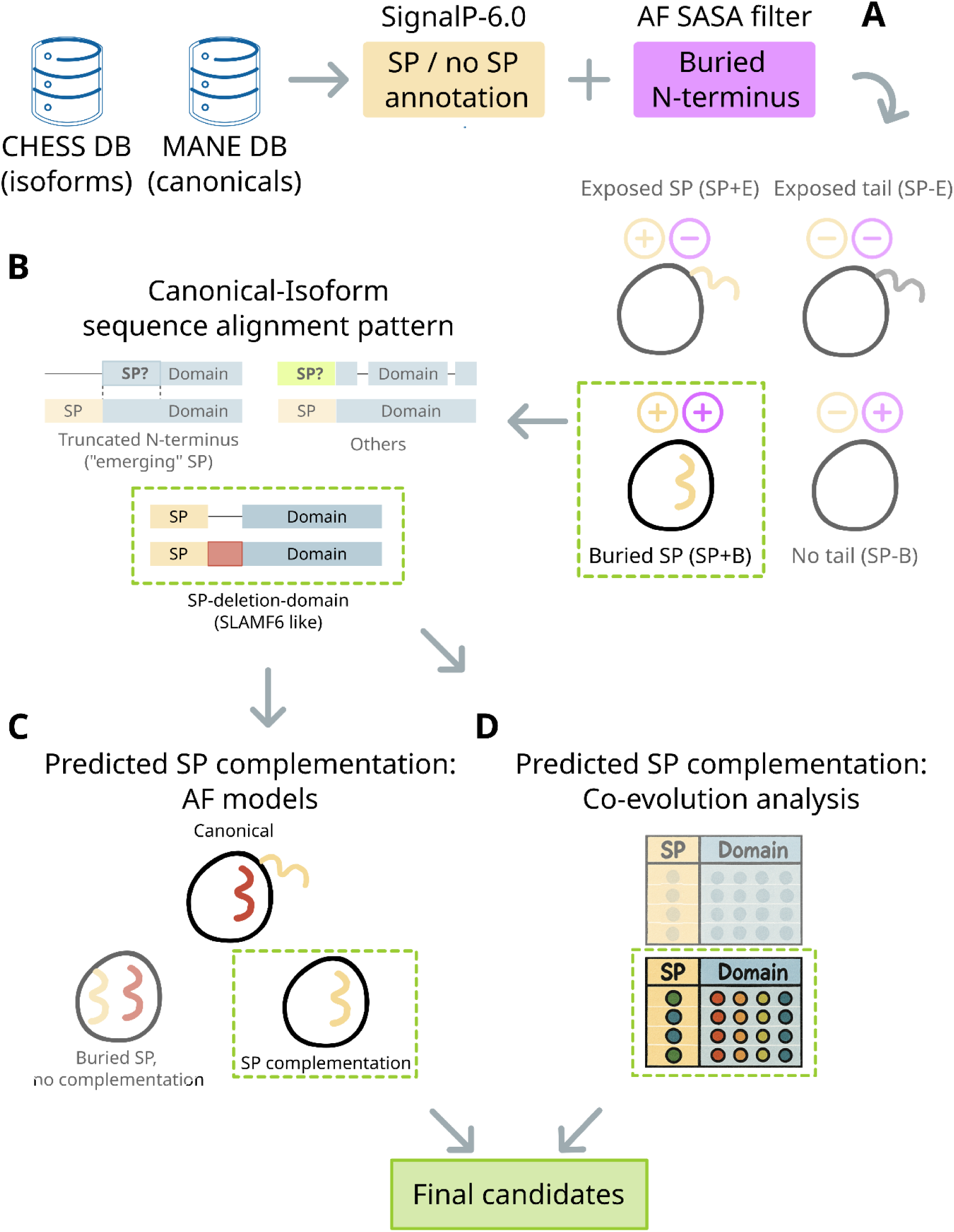
Computational pipeline for detection of SP-complemented human protein isoforms. **(A)** Selection of isoforms with predicted SP (SignalP) and buried (low SASA) n-terminal segment (based on AF model). **(B)** Sequence alignment of selected isoforms to respective canonicals, and selection of isoforms fulfilling pre-defined alignment pattern. **(C)** Structural assessment of SP complementarity to missing segment (based on AF model). **(D)** Co-evolutionary analysis, inferred from the ESM-2 categorical Jacobian.

First, we searched for SPs in all proteins in the dataset. While alternative isoforms have no curated SP annotation (like some canonical proteins do), thanks to recent computational advances the task of SP annotation (prediction) is considered largely solved (55). We used SignalP6 (56) to predict the occurrence and location of SPs for all sequences in CHESS. This resulted in 5,917 MANE proteins and 17,693 isoforms with SP annotation (either confident or suspected, see Methods for annotation cutoffs) **(Figure 3A)**.

Among the isoforms with SP annotation, we wished to detect those in which the SP is structurally involved. For this we used the AF models of the CHESS isoform database (37),(54). For our study, we wished to focus on cases in which the protein contains an SP that is actively involved in stable domain formation. We categorized the models into four classes (**Figure 3A**), based on whether they contain an N-terminal SP (*SP+* and *SP-*, respectively, according to SignalP6 classification), and whether the N-terminus is buried or exposed in the structural model (*B* and *E*, respectively, based on calculated Solvent Accessible Surface Area, SASA; see Methods and **Supplementary Figure S3A**): Class *SP+E* represents canonical proteins translated through the membrane: they contain an exposed SP, since it is usually cleaved and does not contribute further to the protein structure (accordingly, AF models these signal peptides as flanking, exposed and unstructured N-terminal tails, as reported previously (57)). The canonical SLAMF6 isoform represents an example for this class. Classes *SP-B* and *SP-E* in turn represent canonical cytoplasmic proteins: they don’t contain any n-terminal SP, and their retained N-terminus can either be buried and contribute to the structure of the domain (*SP-B*), or an exposed tail (*SP-E*). Class *SP+B* is the group of interest to our study: these proteins do contain an SP, but in contrast to class *SP+E*, this SP is buried and may contribute to domain stabilization (1,598 isoforms, represented by SLAMF6 isoform3). Of note, such examples were previously treated as “AF artifacts” (57). To identify SP complementation cases similar to SLAMF6, we focus hereby on this group of buried SPs.

### Analysis of the “buried SP” group identifies several sequence alignment classes

Not all “buried SP” isoforms follow the same sequence pattern. We aligned the sequence of each isoform in this group to the MANE isoform of the same gene (see Methods). Results of this alignment roughly categorize the members of the “buried SP” group into three classes by alignment pattern (**Figure 3B)**: 1. *SP-deletion-domain*: aligned N-terminal region followed by a deletion, and a second aligned region (124 isoforms). The archetype of this class is SLAMF6 isoform3. 2. *Truncated N-terminus*: Completely missing N-terminal region (including the canonical SP) followed by perfect sequence alignment. This surprising class suggests the emergence of cryptic signal peptides from within the canonical sequence upon splicing, if extracellular location is to be guaranteed (713 isoforms). 3. *Others:* isoforms that do not fit the pattern of these classes. This includes isoforms with new, un-aligned SP sequence, isoforms with negligible changes compared to the canonical sequence, positional mismatches, changes involving predominantly the c-terminal domain, or genes with no annotated MANE isoform (761 isoforms).

We focused on the isoforms in the SP-deletion-domain group (104\124 isoforms, see Methods). While all follow the same alignment pattern and are predicted to bury their SP, only some are predicted to complement the segment missing in the canonical protein. We proceeded to look for potential SP structural complementation (based on AF models, **Figure 3C**) and inferred coevolution (categorical Jacobian, **Figure 3D**).

### Potential signal peptide complementation in several dozen human isoforms

The SP-deletion-domain group includes 104 isoforms (the products of 62 distinct genes) with buried SP and sequence alignment similar to SLAMF6 (**Supplementary Table S1**). Out of 104, for 62 isoforms AF predicts SP complementation (43 genes), *i.e.*, the SP recovers a missing segment (deletion in the canonical sequence). The remaining 42 isoforms (19 genes) in the group are predicted by AF to have a buried SP (not flanking, structured), but no complementation of a missing segment (see **Figure 3C**).

The complementation group is significantly enriched for Ig-like domains as in SLAMF6 (11 isoforms), carbonic anhydrases (3 isoforms) and amyloidogenic glycoprotein e2 domains (2 isoforms) (**Figure 4A, Supplementary Table S2a**); no domain enrichment is detected for the non-complementing group. Significant enrichment for functional interactions is detected in the whole “SP-deletion-domain” group (61 nodes, 21 edges, average node degree: 0.69, p-value: 2.03e-05) and in the complementing group (42 nodes, 12 edges, average node degree: 0.57, p-value: 0.001), but not in the no-complementing group (19 nodes, 1 edge, avg. node degree: 0.11, p-value: 0.3) (**Figure 4B**, **Supplementary Table S2b**).

**Figure 4:**
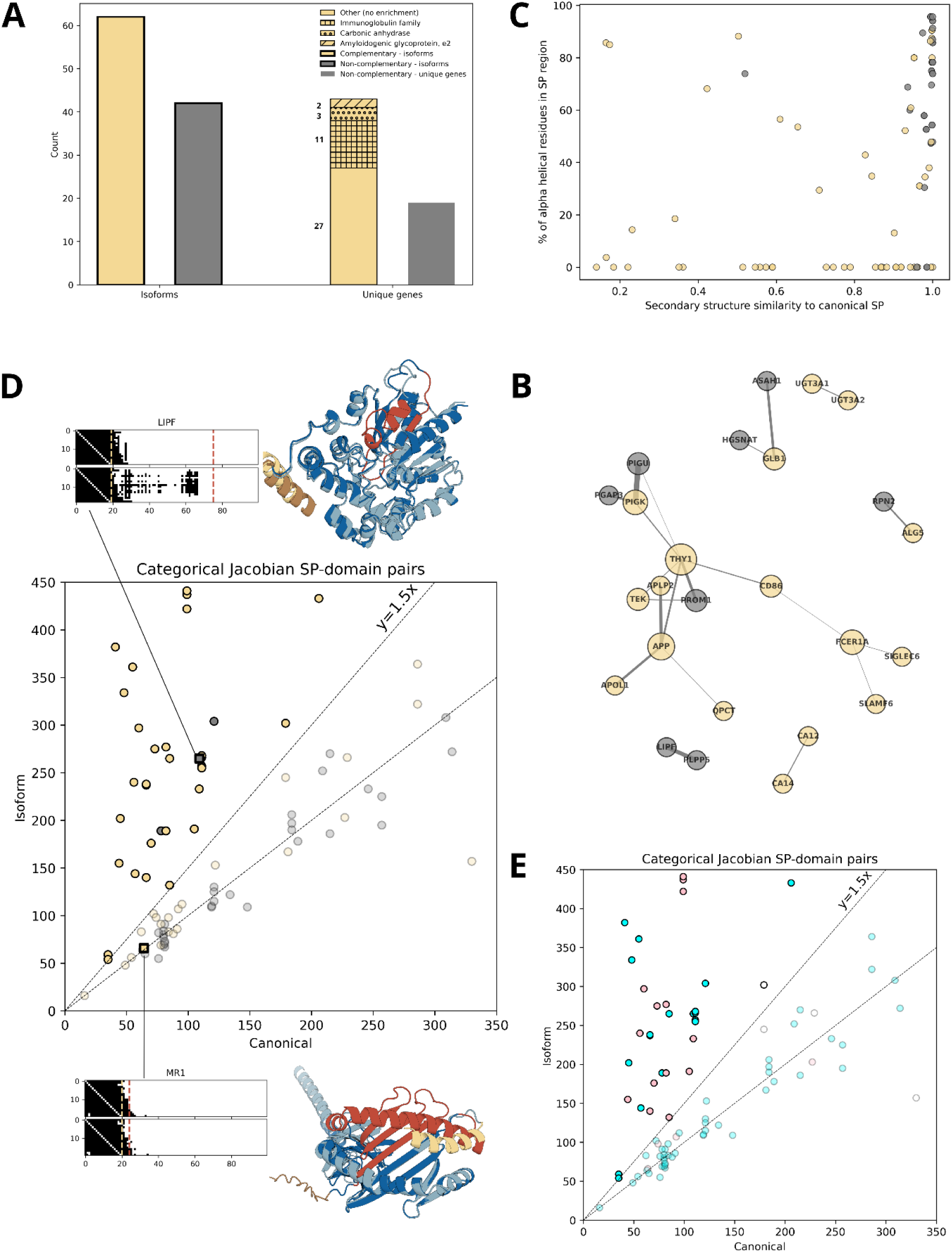
Characterization of potential SP-complementing isoforms. **(A)** Quantitative summary of SP-deletion-domain isoforms (Figure 3C), divided into complementing (wheat) and non-complementing (gray) SP isoforms. The number of isoforms, as well as the number of coding genes that contain such isoforms, is provided. The enrichment of specific domain families in the group of complementing SPs is highlighted as different textures in the barplot (see legend therein, and **Supplementary Table S2a** for details). **(B)** Functional interaction network of complementing and non-complementing group genes. Coloring as in **A** (see **Supplementary Table S2b** for details). **(C)** Secondary structure composition of SP segment with respect to secondary structure similarity of isoform SP region to canonical SP region (cosine similarity of composition vectors). **(D)** Comparison of co-evolutionary signal in SP-domain region between each isoform and its respective canonical counterpart. Coloring as in **A**. Isoforms with number of SP-domain pairs similar to canonical (below y=1.5x diagonal) are transparent. Examples of MR1 (canonical: CHS.3874.17, isoform: CHS.3874.2) and LIPF (canonical: CHS.6738.3, isoform: CHS.6738.4) are highlighted as bold squares connected to additional information. **(E)** Same as panel **D**, colored by deletion in SP segment (cyan) and distal deletion (>5 residues from SP segment; pink).

Complementing SPs are mostly predicted to have β-strand or coiled secondary structure, with a minority of α helices, unlike the mostly α helical non-complementing SPs. Compared to the canonical structures, there is a secondary structure shift in many of the complementing isoforms, *i.e.*, the same sequence is predicted to be helical in the canonical context and non-helical in the isoform context (**Figure 4C**), while no such shift is observed for the non-complementing group. The helix-to-strand shift is supported by the versatile profile of SPs described in the literature (1),(7).

How would the potential of an SP to be buried or structurally complementing affect the confidence of the SignalP prediction? In some examined isoforms the deletion alters the end of the SP segment, which could affect its potential to be cleaved and removed and might also affect the prediction confidence. For the majority of the group (69/104) the predicted SP confidence remains very high (>0.9), and only 27/104 score below 0.5 (**Supplementary Figure S3B)**. However, some promising models are of low SignalP confidence, including the SLAMF6 isoform3 itself (confidence 0.3). Based on the evidence for isoform3 localization, we believe that the low SignalP confidences are suggestive of decreased cleavage ability rather than lack of ability to contribute to membrane translocation. These scores suggest that the examined SPs can potentially moonlight by stabilizing the isoform domains while preserving their localization function.

We proceeded to evaluate the co-evolutionary signal between the SP segments of the complementing isoforms and each complemented domain. Similarly to the analysis performed for SLAMF6 (**Figure 2**), we first calculated the categorical Jacobian contacts for each isoform and canonical sequence. We counted residue pairs predicted to co-evolve where one residue is located in the SP segment, while the other is located in the following domains. Isoforms with a considerable amount of such pairs compared to the respective canonical isoform are considered strong candidates for isoform-specific SP structural complementation (see Methods). 49 isoforms predicted to be SP-complementing by AF show a larger number of co-evolved pairs compared to their respective canonical counterpart, and 34 show 1.5x more pairs (**Figure 4D**). This suggests the existence of a co-evolutionary signal that supports the AF models, and ties the SP to the complemented domain. Importantly, this weaker, less conserved signal “emerges” upon “removal” of the missing segment. Likewise, isoforms predicted by AF to be non-complementing, mainly show no co-evolutionary signal change compared to their canonicals (**Figure 4D**). Curiously, in some of the cases there is a “disagreement” between AF and the evolutionary analysis: for the MR1 isoform, AF predicted a complementation, but no co-evolutionary signal is identified for the isoform sequence (**Figure 4D**, lower panel). Conversely, for the LIPF isoform, AF does not predict an SP complementation, leaving a “hole” where the missing segment used to be in the canonical protein. However, the co-evolutionary signal suggests that the SP may participate in the structure, showing a strong pattern that does not exist for the canonical protein (**Figure 4D**, upper panel). We speculated that isoforms with deletion close in sequence to the SP region are more likely to be complementing (due to smaller structural changes required). Indeed, in a large portion of the final candidates the deletion occurs within 5 residues of the SP, but surprisingly, in many candidates the deletion occurs much further away (**Figure 4E**).

### Examples of strong candidates for SP complementation

To further evaluate the likelihood of our prediction that SPs can provide structural complementation to stabilize truncated domains in an isoform, we characterized in more detail the set of strong candidates, *i.e.*, isoforms with high SP complementing potential both based on their AF structural model, as well as their categorical Jacobian. We inspected two additional potential lines of evidence: *1. Detection of SP segments in our candidate isoforms by Mass Spectrometry (MS).* It is known that in general SPs are less likely to be detected in MS experiments due to their usual short lifetime (cleavage followed by degradation) (58),(59). Moreover, hydrophobicity and limitations of MS methods (regarding protein termini) further reduce their probability to be detected (60),(61). We therefore assume that MS detection of SPs could reflect their continued presence, and therefore functional importance. *2. Detection of conserved sequence motifs.* SP complementation is more likely to occur with sequence motifs conserved between the missing segments and their complementing SPs. We present here three final candidates with compelling support by MS data (see Methods), sequence complementarity or both, alongside distinct predicted co-evolutionary signals. These examples represent the enriched domains (**Figure 4A)**, namely immunoglobulin like (THY1, FCER1A) and carbonic anhydrases (CA12) domains.

THY1 is a single domain membrane glycoprotein, predicted by AF to form an Immunoglobulin-V like structure composed of 2 β sheets of 4 strands each (**Figure 5A**). The α helical flanking SP is cleaved and does not contribute to the structure of the domain. Our analysis detected isoform CHS.10184.18 in which the first 12 residues are aligned, followed by a deletion (V13-L29), and a second aligned region. The deletion, starting immediately after the SP segment, is predicted to remove the first β-strand (n-terminal) and likely alters the SP cleavage site (predicted at position 19). The isoform structure is predicted to be nearly identical to the canonical protein, with the previously helical signal peptide fully recovering the missing β strand. Categorical Jacobian analysis clearly shows new residue pairs in the isoform compared to the canonical protein. These pairs are in agreement with the AF model, “pinning” the SP into the first β strand position (**Figure 5A**). Strikingly, several proteomic MS experiments have detected one consecutive peptide of the isoform sequence that covers residues M1-R39 and includes the newly created joint (positions T12-V13, **Figure 5A**). This experimental evidence suggests that 1) the isoform exists at the protein level, and 2) the signal peptide is an integral part of the isoform structure.

**Figure 5:**
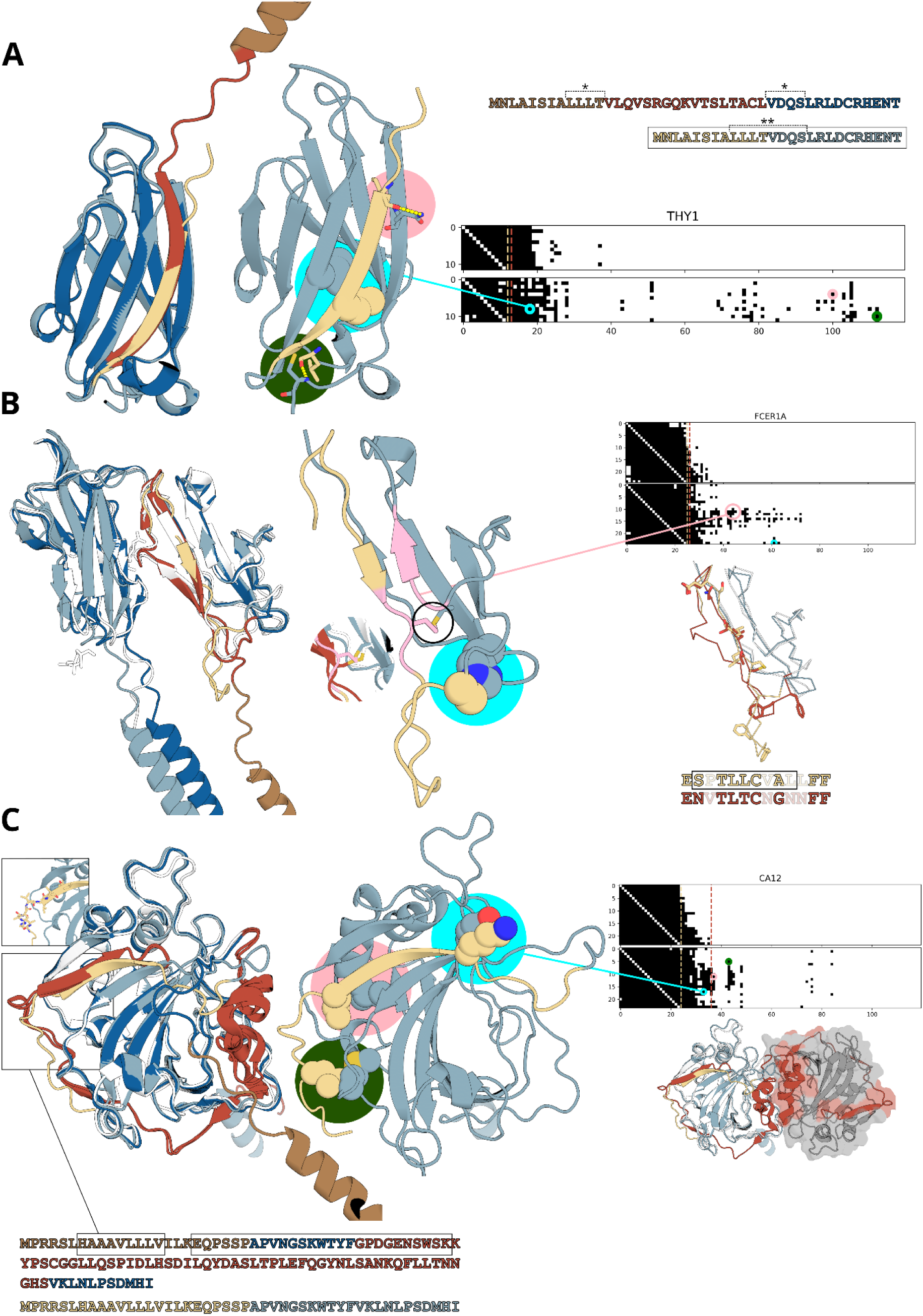
Examples of isoforms with truncated domains that are complemented by an SP. **(A)** Thy1 (CHS.10184.5, CHS.10184.18). **(B)** FC receptor of IgE (CHS.3426.6, CHS.3426.5). **(C)** Carbonic anhydrase (CHS.17547.7, CHS.17547.5). Overlay of canonical and isoform models (left), isoform model (middle), categorical Jacobian (right). Color scheme as in Figure 1 (White - crystal structures, dark colors - canonical isoform, light colors - alternative isoform). N-terminal canonical and isoform sequences colored accordingly. MS detected sequences are delineated by boxes. The SP-domain part of categorical Jacobian is shown (canonical - upper, isoform - lower), with representative residue pairs marked with colored circles, corresponding to the same color circle highlights of the 3D structures. **(A) THY1** - isoform sequence detected by MS delineated by box; segment proximal and distal to the deletion in the canonical sequence delineated with dashes and *, new joint in isoform sequence marked with **. **(B) FCER1A -** Middle - isoform model, zoomed in on complementation area; S-S bridge complemented by SP Cys 12 marked by black circle, and shown in details compared to crystal structure (white, dark red). Right - complementing SP overlay with crystal structure deletion area shown in ribbon, identical/similar residues shown in sticks, and highlighted in sequence snippet. **(C) CA12 -** Left - homodimer interface shown in cartoon, with gray transparent surface, deletion area in dark red. MS detected segments shown in boxes (corresponding to box highlights of 3D structure).

The high affinity immunoglobulin epsilon receptor subunit α (FCER1A) is a well-studied FC receptor of IgE. Its structure is composed of 2 adjacent immunoglobulin-like domains, with β sheets of 3-4 strands (**Figure 5B**) (8K7R (62)). High resolution solved structures of the canonical protein do not include the SP, since it is cleaved and not involved in the structure (the AF model of the canonical protein predicts a flanking, helical SP that does not interact with the domain structure). Isoform CHS.3426.5 is missing a segment (V26-E58) that starts immediately after the SP segment, predicted to remove the first three β-strands and a following loop. The isoform structure is predicted to use the SP to replace two out of the three missing β strands, but in contrast to the SLAMF6 isoform discussed above (**Figure 1**), the missing segment in the IgE FC receptor isoform is not located in the region that forms the interface with its IgE ligand. Categorical Jacobian analysis reveals a strong co-evolution signal, in agreement with the AF model, locating the SP β strand to complement the truncated β sheet. The complementation segment shows an impressive sequence similarity (**Figure 5B**), with overall good structural alignment of the similar positions, except for the loop (including the sequence F56,F57 aligned to isoform F17,F18) that is not structurally aligned. Furthermore, in the canonical protein an S-S bridge is created between residues C51-C93. C12 in the SP replaces C51 in the missing segment to recapitulate this S-S bridge (**Figure 5B**). Promisingly, most of this complementing segment is also detected by MS (**Figure 5B**).

Carbonic Anhydrase 12 (CA12) is a member of a large family of enzymes performing hydration of carbon dioxide. It adopts a carbonic anhydrase fold, composed of a main β sheet of 10 strands, surrounded by several loops and α helices (**Figure 5C**) (PDB id 6R71 (63)). As expected, the many solved structures of CA12 do not include the canonical SP, since it is cleaved and not involved (modeled as a detached helical fragment by AF). Isoform CHS.17547.5 lacks a long segment (G36-S95), but is otherwise identical in sequence. The missing segment is predicted to remove a long coiled/helical stretch located in the homo-oligomerization interface of the canonical protein (**Figure 5C**). Additionally, the deletion removes one of the β strands composing the main β sheet. The isoform SP is predicted to recover the missing β strand along-side a short, coiled stretch. As with previous examples, co-evolutionary signals support the AF model in several sites across the complementation. While no clear sequence similarity pattern is shared between the missing and the complementing segments, a large portion of the complementing segment is detected by MS (**Figure 5C**), suggesting its presence in the protein.

## Discussion

Signal peptides play a crucial role for transmembrane and extracellular protein translation, but are usually cleaved, and have been mostly considered irrelevant for the structure of the mature protein that they help localize. In this work, we explored the potential of human SPs to play a significant role in protein stabilization, with implications for function. The sequence and structural versatility of SPs has been long established, but their “moonlighting” roles aside from localization are only starting to be discovered. We present a new perspective, suggesting that SP sequences may have “context dependent” roles, changing between canonical and alternatively spliced proteins. We hypothesize that segments missing from splice isoforms (compared to canonical proteins) could be “replaced” or complemented by SPs. We provide in-silico predictions that support this hypothesis alongside experimental evidence, proposing several dozen protein isoforms where SP complementation may occur. Complementing isoforms are enriched for immunoglobulin and carbonic anhydrase domains, and are non-trivially enriched for functional interactions, suggesting a possible broader relationship.

This perspective suggests that some proteins harbor “replacement parts” in the form of signal peptides, which are structurally “recruited” upon specific splicing events. Previous work that explored AF models of proteins with SPs, stumbled upon a minority of SPs predicted to be buried, which they deemed to be AF artifacts (57). We provide biological context in which such predictions make structural and functional sense: for some, e.g SLAMF6 and FCER1A, we demonstrate surprising sequence similarity to a missing domain segment, suggesting that the SP acts as a “spare tire”; Others such as THY1 show strong MS evidence suggesting that the SP is unlikely to be cleaved, and likely to be involved in the mature protein structure.

In general, AF predictions have been thoroughly validated, and could even be used to refine experimentally solved structures. However, they are not free of mistakes. These mistakes or biases are thought to be especially strong for protein isoforms, given highly similar sequence to canonical sequence, lack of sufficient MSA depth or MSA misalignments of short regions, failure to model fold changes, lack of representation in structural training data and more. A major limitation of this work is the reliance on AF models, to model an inherently difficult structural biology question. However, we are encouraged to believe that our predictions are unlikely to be due to AF biases. This is reflected by the structural models in support of our hypothesis for SLAMF6 using methods that are not based on deep learning or MSAs, as well as by the unexpected signal predicted/detected by the ESM categorical Jacobian that connects SP regions to complemented domain regions (without the direct involvement of MSA or structural information). We showcase “copied” sub-folds predicted by AF, which in isolation seem biophysically unlikely, but could be easily accounted for in the context of oligomerization. Moreover, while AF may predict identical sequence segments as structural components “memorized” from larger (e.g. canonical) proteins, signal peptide structures are not available in the PDB; making it more likely that complementing SPs “fit” in their predicted conformations.

The accumulation of evidence in support of SP complementation from different predictions, is sufficient to suggest an actual biological phenomenon, rather than random noise. Our hypothesis builds on numerous pillars, each of which will require further experimental validation: (1) What is the biological relevance of the studied isoforms? (2) Are the isoforms indeed located as predicted based on SP annotations? (3) Do we actually see effects on stability and reduced degradation, as suggested from MS detection experiments?; and (4) What is the actual role of this structural complementation by SPs? How does it influence observed functional effects (e.g. for SLAMF6)? Future studies should test the stability of the candidate isoforms with respect to the predicted SP structure, and the effects of destabilized SP complemented isoforms on the phenotype of the relevant genes.

Signal peptide complementation will have an impact on numerous areas of the study of proteins. Nearly one third of human proteins are known to include a signal peptide, alongside many more in other species. It is custom to remove the SP region for protein expression, or to replace native SPs with optimized SP sequences, under the assumption that the mature protein is unaffected. Our perspective suggests that these procedures might dramatically change the protein structure or function. Moreover, SPs from thousands of proteins are cleaved off and are free to engage in peptide-protein interactions. Here we demonstrate the potential of these peptides to stabilize the same protein they localize, but potentially, this could be extended to any other protein engaging with a soluble peptide. Finally, for splice isoforms, many questions arise - how does SP complementation allow to diversify the protein repertoire? What new interactions are created by the similar but not identical domains/interfaces complemented by SPs? The structural significance of SPs remains a fertile ground for future exploration that we expect will lead to new research frontiers.

## Methods

### Canonical and alternative splicing isoform databases

All isoform sequences and ID mappings were retrieved from the freely available <u>isoform.io</u> DB v1.3 (54) based on (CHESS 3) (37). Annotation of canonical isoforms was performed by matching MANE DB v1.4 (38) sequences to CHESS isoform sequences. CHESS isoforms matching any MANE class (MANE, MANE_select, *etc*), were considered MANE canonical isoforms. A total of 237,275 CHESS isoforms were processed, among them 33,851 annotated as MANE isoforms.

### Signal peptide annotation using SignalP-6.0

Annotation of SP regions and confidence was performed using SignalP-6.0 (56), local version, with ‘slow_sequential’ model and default parameters. A total of 23,880 SP annotated sequences were considered: any sequence (isoform as well as MANE) with SignalP prediction confidence of 0.5 and above was considered ‘confident SP’ (19,668 sequences), and predictions with confidence 0.1 - 0.5 were considered ‘possible SP’ (4,212 sequences); all other sequences, the vast majority, were discarded (213,395). Prediction probabilities of the aforementioned groups are described in **Supplementary Figure S4**. For the ‘possible SP’ group, SP boundaries are missing (not automatically predicted by SignalP), hence the end position was assigned at residue 23, based on average length of signal peptides in the confident set.

### Detection of buried SP isoforms

Structures of isoforms annotated as “confident SP” or “possible SP” were analyzed for the burial of the SP. The solvent accessible surface area (SASA) of the first (N-terminal) 100 residues was computed using the Biopython package PDB.SASA module (64), and used as a rough approximation of buried *vs* exposed areas. SASA was computed using the default parameters (Shrake-Rupley algorithm (65)), at the residue level (level = ‘R’ parameter) and validated to match the full structure SASA (level ‘S’). This resulted in a vector of length 100 of by-residue SASA values. The vector was split into two: the SP segment according to SignalP annotation was compared to the following segment. For isoforms in the “possible SP” group, the SP segment was defined 1:23, as explained above (e.g. [1:23] vs [24:100]. Average SASA value for each segment was computed. Isoforms with average SP SASA ≤ 90Å² were considered “buried SP”. This SASA cutoff was defined by comparison to average values of folded domains in the data set (see **Supplementary Figure S3A**).

### Sequence pattern detection

Sequence alignment of splice isoforms to their respective MANE (canonical) isoform was performed using the Align module of Biopython (64) v1.84 with parameters (manually adjusted): open_gap_score = -5, extend_gap_score = -0.1, target_end_gap_score = 0.0, query_end_gap_score = 0.0. Isoforms were classified into groups based on their alignment pattern: 1. Identical to MANE; 2. No MANE assigned for gene; 3. Aligned SP with deletion pattern (pattern of SLAMF6 isoform3); 4. Full n-terminal truncation with perfect subsequent alignment; and 5. Others: alignments not following previously described patterns. Alignment patterns were defined using the following regex strings: SP-and-deletion = r’\|{5,}\.?-{5,}\.?\|{5,}’; n-terminal truncation = r’-{5,}\.?\|{5,}’.

### Structural detection of SP complementation

Each isoform in the SP-and-deletion set was structurally aligned to its annotated canonical (MANE). For structural alignment we used the command “super” from the Pymol-open-source package v3.0.0 (66) in 3 separate rounds (full protein, 100 n-terminal residues, 50 n-terminal residues) to account for possible local alignments of the N-terminal domain and SP. Using the best alignment, Root Mean Square Deviation (RMSD) was calculated between the isoform SP segment and the deletion segment of the canonical protein. Since the complementation could be partial, the distance between each Cα atom of the SP segment to each Cα atom of the deletion segment is computed, and the minimal distance is retained. Isoforms with minimal distance within 2.5Å were classified as “complementing SPs”, while others as “non complementing” SPs (**Figure S3C)**.

### Categorical Jacobian analysis

Categorical Jacobian analysis was performed using code by Zhang *et al.* (47) (https://github.com/zzhangzzhang/pLMs-interpretability), locally adapted for SLURM on a GPU cluster. Default parameters and functions were used. A permissive cutoff of top 15xL(protein length) predicted pairs were selected for each Jacobian contact matrix to allow for detection of subtle signals, similar to (46) who used 15L/2 (best of 3 different coevolution analysis methods). For a protein of length L=300, top 15L pairs correspond roughly to top 10% interactions and for L=1000, roughly to top 3%. Residue pairs showcased in **Figure 2** and **5** were selected by a cutoff of 5 Å heavy atom distance, similar to (67).

<u>Evaluation of SP-domain coevolution pairs:</u> of the top 15L pairs in the Jacobian contact matrix, SP-domain interaction pairs (i,j) were counted such that residue i is within the SP segment boundaries ([1-SP_end]) and residue j is outside the SP segment boundaries. The part of SP segment identical between each canonical and respective isoform (termed “common SP”) was used to define boundaries for this analysis: SP boundaries for canonical SLAMF6 e.g. are defined 1:21; for isoform3, first 17 residues are identical compared, while the rest are removed. Hence only the first 17 residues were used for Jacobian analysis for both canonical and isoform sequences ([1-SP_end] was defined [1-17]). “common SP” values for SP-deletion-domain pairs are reported in **Supplementary Table S1**.

Deletion in SP segment *vs* distal deletion (**Figure 4E**): isoforms with a deletion starting within the SP boundaries and up to +5 residues of SP_end were defined as “Deletion in SP segment” (otherwise - “distal deletion”).

### Secondary structure annotation and comparison

Secondary structure annotation was performed using the pydssp package https://github.com/ShintaroMinami/PyDSSP, a python implementation of DSSP (68). Each SP secondary structure was summarized as a percentage vector of each SS2 element (e.g., 0.3 H, 0.1 E, 0.6 C). Secondary structure similarity between SP segments of isoforms and their respective canonical proteins were computed as the cosine similarity between the canonical and the isoform H,E,C summary vector (see **Supplementary Table S1**).

### Domain and functional interaction enrichment analysis

104 Isoforms of the SP-and-deletion group (**Figures 3C-D and 4, Supplementary Table S1**) were analyzed for domain and functional interaction enrichment using the STRING v.12 online web server (https://string-db.org (69)). Default parameters were used with no addition of interactors outside the input group. Enrichment values are reported in **Supplementary Table S2a,b**.

### Mass spectrometry data

All MS coverage data was extracted from the PeptideAtlas project web server (https://peptideatlas.org/ (70)). The following query was used: https://db.systemsbiology.net/sbeams/cgi/PeptideAtlas/GetProtein?atlas_build_id=592&protein_name=XXX&action=QUERY), where XXX is the Uniprot ID in the following table:

| Protein | CHESS identifier | Uniprot ID |
| --- | --- | --- |
| THY1 | CHS.10184.18 | J3QRJ3 |
| FCER1A | CHS.3426.5 | E9PRN1 |
| CA12 | CHS.17547.5 | B3KUB4 |

### Structural search of protein isoform models

Isoform analysis data and structural models were retrieved from https://github.com/HWaymentSteele/Isoforms_benchmark_2024 (47). AF2 models of MYG and CASP9 were searched against the PDB using Foldseek (71) with default parameters. Search hits were screened and analyzed manually.

### Rosetta and I-TASSER models of SLAMF6 isoform3

The Rosetta (52) model of SLAMF6 isoform3 v-domain was predicted with a local version of Rosetta, linux release, v.2018.09.60072, using the *ab-initio* relax protocol with default parameters, creating n=200,000 decoys and no use of templates. The model presented in **Figure 1** is the top 6 ranked model by Rosetta score (cluster representative).

Standard (non deep learning) I-TASSER models were predicted using the webserver (https://zhanggroup.org/I-TASSER/ (53)) with default parameters and templates.

## Code Availability

All code used to perform the described analyses are available in https://github.com/Furman-Lab/Signal_peptide_modeling.

## Data Availability

Isoform sequences and AlphaFold models are freely available in CHESS https://ccb.jhu.edu/chess/ and https://www.isoform.io/home.

All data generated in this study is provided in https://github.com/Furman-Lab/Signal_peptide_modeling.

## Author Contributions

T.T. and O.S-F. conceived the study. T.T. collected data, designed and performed data analysis with supervision of O.S-F. Both authors wrote and edited the manuscript.

## AI usage declaration

Large language models such as Claude and ChatGPT were used to refine code written by T.T. All text in the manuscript was written by T.T. and O.S-F. The illustration in **Figure 2A** was manually drawn by T.T. and aesthetically refined using ChatGPT.

## Acknowledgments

This work was supported, in whole or in part, by the Israel Science Foundation, founded by the Israel Academy of Science and Humanities (grant numbers 717/2017, 301/2021 to O.S.F.) and by the Clore Foundation Scholarship for PhD students (T.T.).

## Supplementary Materials

**Figure S1:**
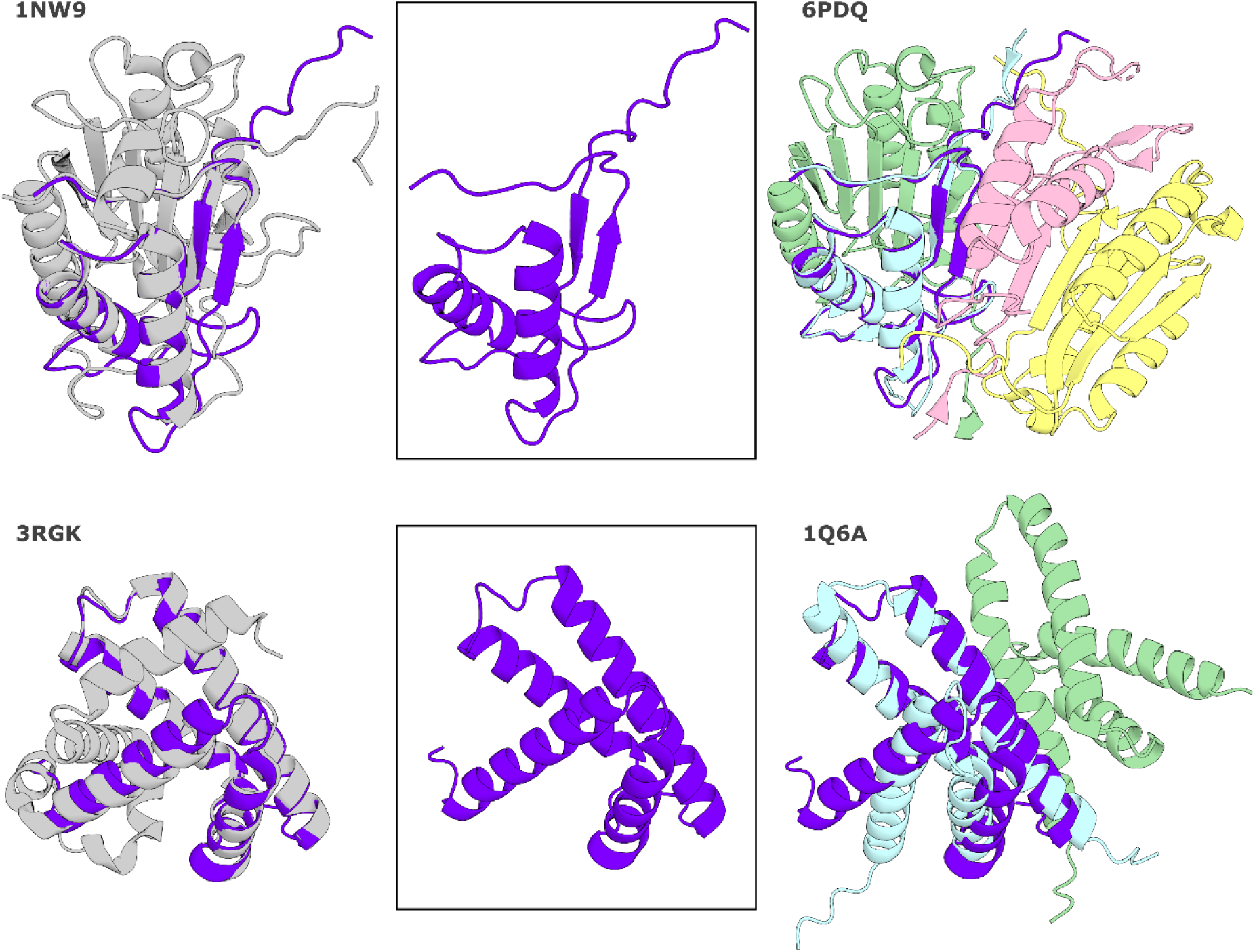
AF2 models of isoforms of CAS9 (upper) and MYG (lower) examined by Zhang *et al.* (*47*) may be well explained in the context of larger protein complexes and potentially fold as predicted. Middle: Splice isoform model (purple); Left: Canonical isoform (gray), with splice isoform model superimposed; Right: Isoform superimposed onto structure of complex to highlight potential structural complementation (matched chain in cyan). Crystal structure PDB ids: CASP9: 1NW9 (72) and 6PDQ (73); MYG - 3RGK (74) and 1Q6A (75).

**Figure S2:**
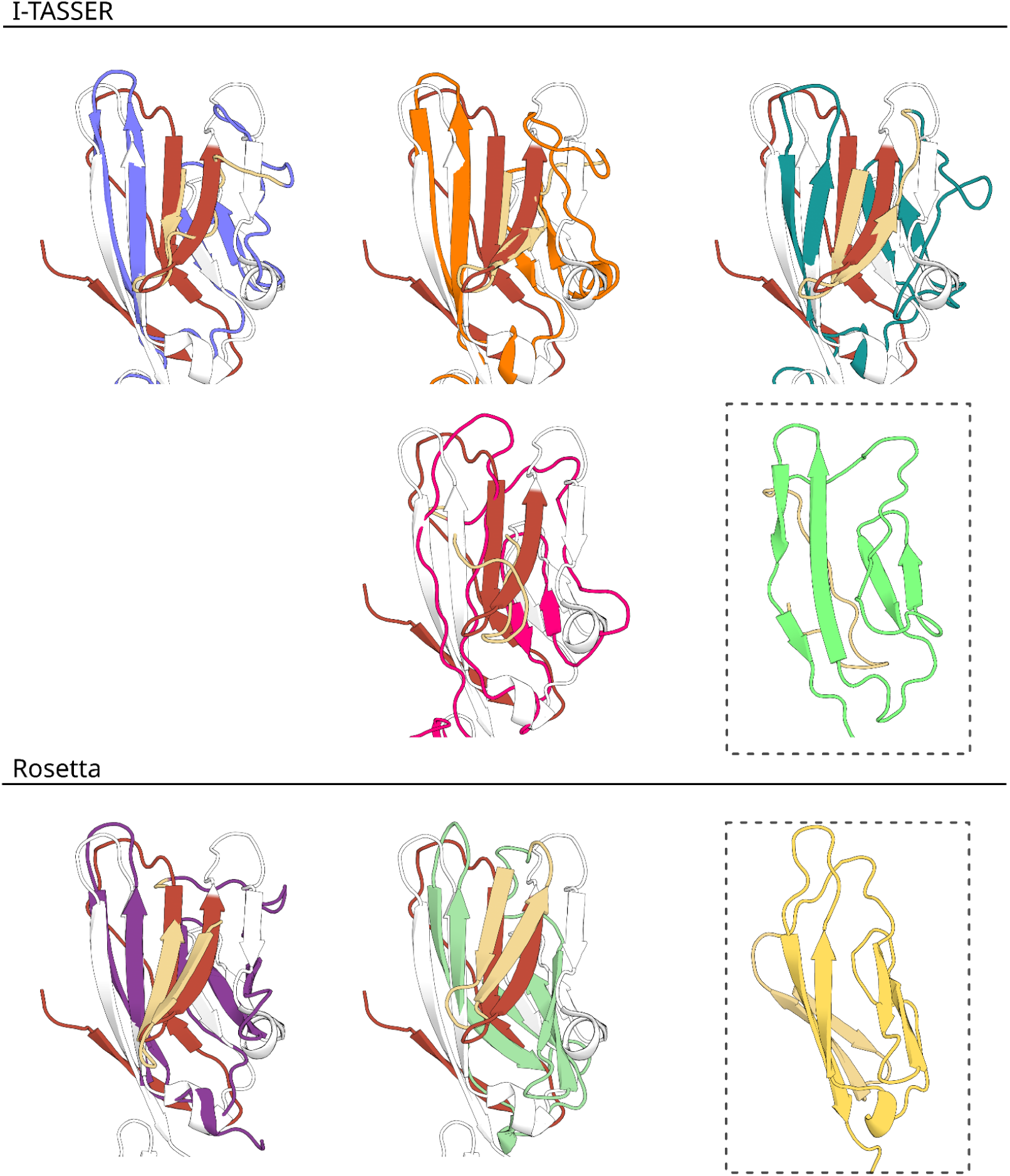
Structural models of isoforms support complementation by signal peptides. I-TASSER models of SLAMF6 isoform3 are similar to corresponding models generated using AF2 as well as Rosetta *ab initio*. Both methods converge into SP complementing models of the front β-sheet face (strands C,C’; models superimposed onto crystal structure), with some models complementing the posterior face of the β sheet (dashed boxes, no crystal structure shown). Color scheme as in Figure 1.

**Figure S3:**
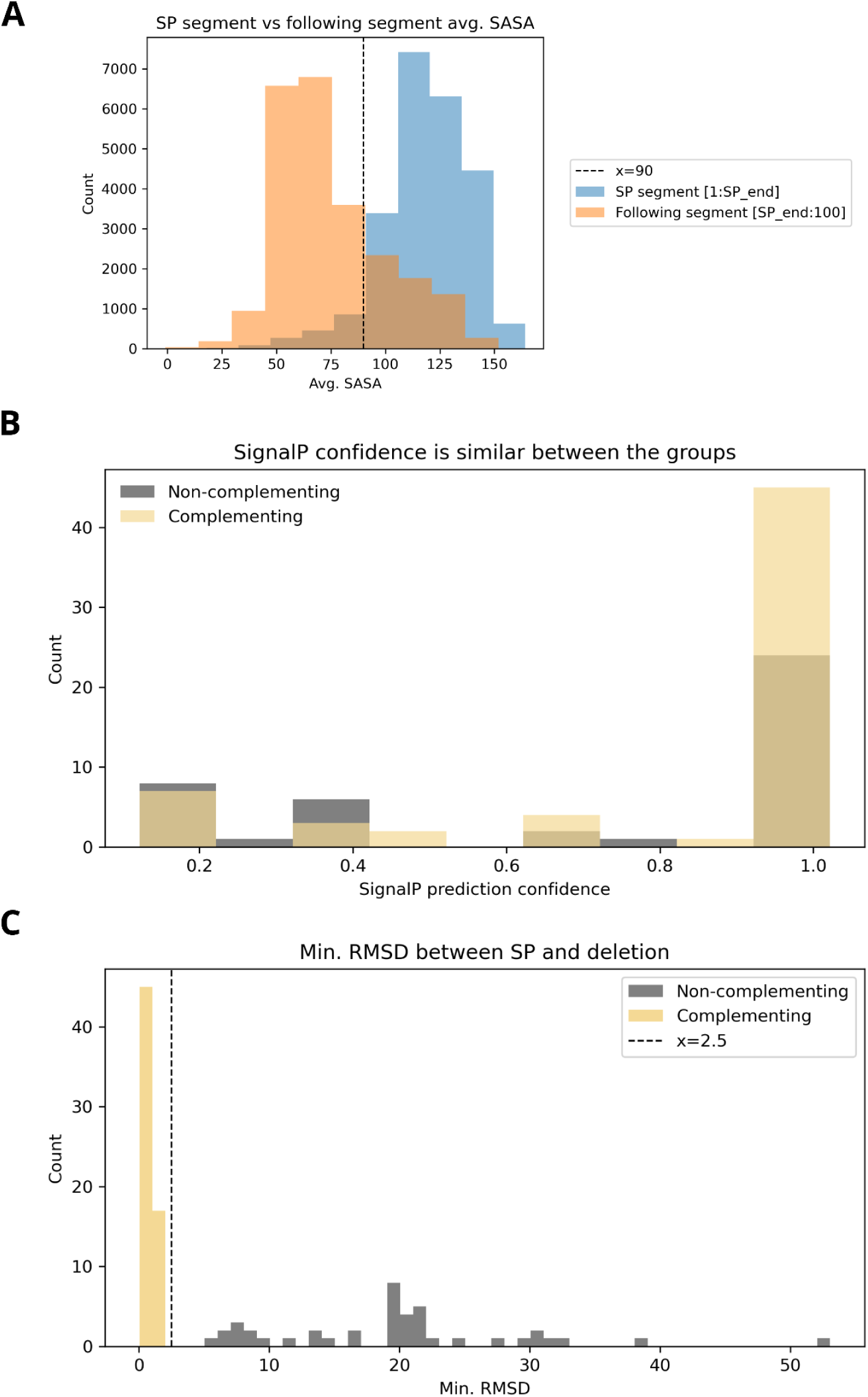
Characteristic features of complementing SPs, in comparison to their following sequence (A), as well as to non-complementing SPs (B,C). **(A)** Average Solvent Accessible Surface Area (SASA) values of N-terminal segments compared to following segments ([1:SP_end] vs [SP_end:100]) in AF2 models of the analyzed dataset. A cutoff of 90Å^2^ average SASA of SP residues (dashed vertical line) was used to define buried SPs. **(B)** Histogram of SignalP prediction confidence values for complementing and non-complementing SP groups (wheat and gray, respectively). Values are overall high, and similar between the groups. **(C)** Minimal RMSD between SP segment and deletion segment.

**Figure S4:**
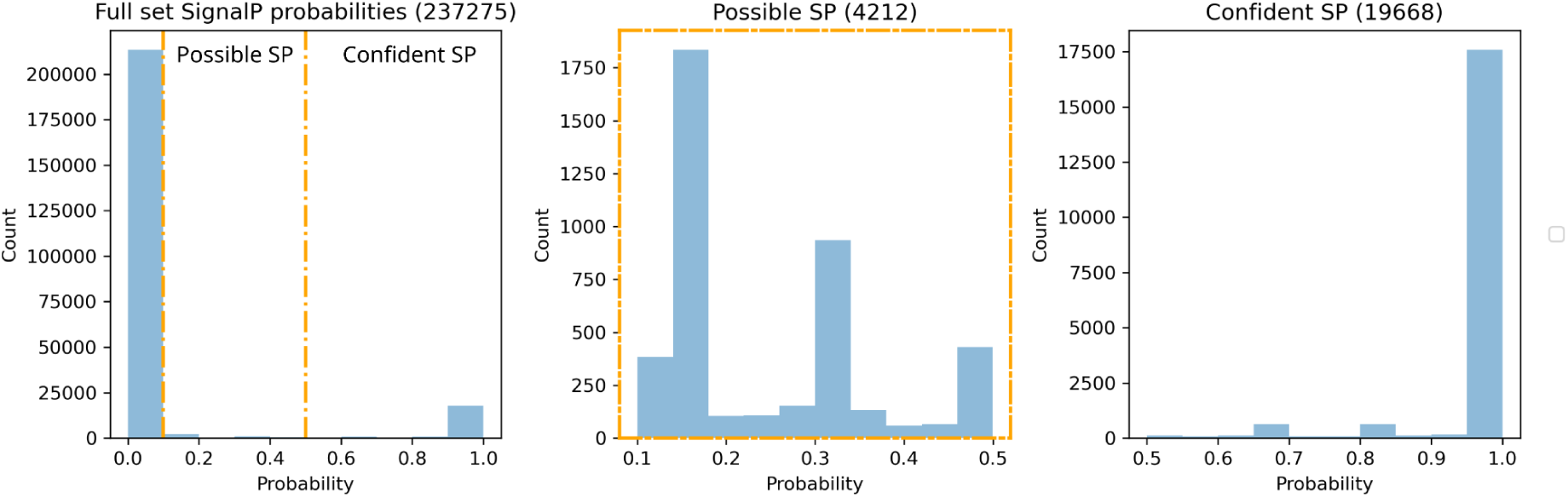
Histogram of SignalP prediction confidence values for the whole data (left), marking the “possible SP” group (orange dashed lines), and “confident SP” group. Zoomed in histograms for possible SP and confident SP shown middle and right respectively.

**Table S1: Summary table of the SP-deletion-domain group, 104 isoforms (**see Figure 4). For each entry, details about the canonical and alternative isoform are provided. This includes the range and sequence of the SP complementation, as well as structural similarity. Provided as tsv file.

**Table S2: Functional interactions between SP-deletion-domain genes** (see Figure 4A**,B**). **(A)** Domain enrichment within the group as predicted by STRING. **(B)** Functional associations (edges) between genes in the group (nodes) and the confidence score as predicted by STRING. Provided as tsv file.

